# Arousal-State Dependent Alterations in VTA-GABAergic Neural Activity

**DOI:** 10.1101/770313

**Authors:** Ada Eban-Rothschild, Jeremy C. Borniger, Gideon Rothschild, William J. Giardino, Joshua G. Morrow, Luis de Lecea

## Abstract

Decades of research have implicated the ventral tegmental area (VTA) in motivation, reinforcement learning and reward processing. We and others recently demonstrated that it also serves as an important node in sleep/wake circuitry. Specifically, VTA-dopaminergic neuron activation is sufficient to drive wakefulness and necessary for the maintenance of wakefulness. However, the role of VTA gamma-aminobutyric acid (GABA)-expressing neurons in arousal regulation is not fully understood. It is still unclear whether VTA-GABAergic neurons predictably alter their firing properties across arousal states, what is the nature of interactions between VTA-GABAergic activity and cortical neural oscillations, and how activity in VTA-GABAergic neurons relates to VTA-dopaminergic neurons in the context of sleep/wake regulation. To address these questions, we simultaneously recorded population activity from VTA-GABAergic or VTA-dopaminergic neurons and EEG/EMG signals during spontaneous sleep/wake states and in the presence of salient stimuli in freely-behaving male mice. We observed that VTA-GABAergic neurons exhibit robust arousal-state-dependent alterations in population activity, with high activity and calcium transients during wakefulness and rapid-eye-movement (REM) sleep compared to non-REM (NREM) sleep. During wakefulness, population activity of VTA-GABAergic neurons, but not VTA-dopaminergic neurons, was positively correlated with EEG gamma power and negatively correlated with EEG theta power. During NREM sleep, population activity in both VTA-GABAergic and VTA-dopaminergic neurons negatively correlated with delta, theta, and sigma EEG power bands. Salient stimuli, with both positive and negative valence, activated VTA-GABAergic neurons. The strongest activation was observed for social stimuli irrespective of valence. Together, our data indicate that VTA-GABAergic neurons, like their dopaminergic counterparts, drastically alter their activity across sleep-wake states. Changes in their activity predicts cortical oscillatory patterns reflected in the EEG, which are distinct from EEG spectra associated with dopaminergic neural activity.

**Statement of Significance:** Little is known about how ventral tegmental area (VTA) neural ensembles couple arousal to motivated behaviors. Using cell-type specific genetic tools, we investigated the population activity of GABAergic and dopaminergic neurons within the VTA across sleep/wake states and in the presence of salient stimuli. We demonstrate that coordinated neural activity within VTA-GABAergic neurons peaks during wakefulness and REM sleep. Furthermore, neuronal activity in VTA-GABAergic neurons is correlated with high frequency, low amplitude cortical oscillations during waking, but negatively correlated with high amplitude slower frequency oscillations during NREM sleep. Our results demonstrate that VTA-GABAergic neuronal activity is tightly linked to cortical arousal and highlight this population as a potential important node in sleep/wake regulation.

## Introduction

Motivated behaviors are fundamentally linked to arousal, however, little is known about the neural mechanisms that couple motivational processes with sleep-wake regulation (Eban-Rothschild et al., 2018). The ventral tegmental area (VTA), which is comprised of 55-65% dopaminergic, 35-40% gamma-aminobutyric acid (GABA) and ∼2% glutamatergic neurons (Nair-Roberts et al., 2008; Morales and Margolis, 2017), is a key brain structure in the regulation of motivation, reinforcement learning and reward processing (Berridge, 2007; Salamone and Correa, 2012; Berke, 2018). VTA-GABAergic neurons have been suggested to tightly control activity in intermingled VTA-dopaminergic neurons in the context of motivation and reward (Cohen et al., 2012; van Zessen et al., 2012), yet their regulatory role in the context of sleep and wakefulness is not fully understood.

Neuronal activity in the VTA exhibits arousal-state-dependent alterations, with increased activity during wakefulness and REM sleep compared to NREM sleep (Miller et al., 1983; Fifel et al., 2018). Although initial studies suggested that VTA-dopaminergic neurons do not change their mean firing rate across sleep/wake states (Miller et al., 1983; Steinfels et al., 1983), more recent examinations revealed marked arousal state-dependent alterations in both firing rate and firing pattern (Dahan et al., 2007; Gomperts et al., 2015). Moreover, using cell-type specific fiber photometry recordings, we demonstrated that at the population level, VTA-dopaminergic neurons show strong arousal-state-dependent alterations in activity (Eban-Rothschild et al., 2016). Early single-unit electrophysiology recordings in rats suggested that activity in VTA-GABAergic neurons varies significantly across arousal state, with high firing rates during active waking and REM sleep (Lee et al., 2001). An additional study demonstrated activation, via cFos immunolabeling, of VTA-GABAergic neurons following recovery from REM sleep deprivation (Maloney et al., 2002)–suggesting that these neurons influence or monitor discrete arousal states. Nonetheless, there is a great challenge to accurately differentiate between different cell types in freely moving wild type animals and conventional electrophysiological characteristics used to differentiate between VTA cell types has been debated (Margolis et al., 2006; Lammel et al., 2008; Ungless and Grace, 2012). Moreover, it is still unclear whether activity in VTA-GABAergic neurons precedes or lags behind arousal state transitions, how population activity in the VTA is related to different components of the cortical EEG and how VTA-GABAergic population dynamics relate to that of co-mingled dopaminergic neurons. A few recent studies have shed some light onto these questions, demonstrating that VTA-GABAergic neurons bidirectionally regulate wakefulness in part via their projections to the lateral hypothalamus (LH) (Takata et al., 2018; Chowdhury et al., 2019; Yu et al., 2019b). However, there are some critical discrepancies between these studies, as one suggests that VTA-GABAergic neurons are predominantly wake and REM sleep active (Yu et al., 2019), while the other provides evidence that these neurons fire mostly during NREM sleep (Chowdhury et al., 2019). Therefore, the exact phenotype of VTA-GABAergic neurons across arousal states remains unclear.

To address these questions, we used Cre-dependent viral vectors to express the calcium indicator GCaMP6 exclusively in either VTA-GABAergic or VTA-dopaminergic neurons. We then used fiber photometry in tandem with electroencephalography/electromyography (EEG/EMG) recordings to monitor spontaneous arousal-state dependent changes in population activity. We also recorded calcium and EEG measures during a battery of behavioral challenges (social and non-social).

## Methods

### Animals

We used adult (>8 wks old) male vesicular GABA transporter (Vgat)-ires-Cre (Jackson Labs stock #028862) knock-in mice backcrossed on the C57BL/6J wild-type background (for more than 10 generations) and tyrosine hydroxylase (Th)-IRES-Cre mice obtained from the European Mouse Mutant Archive and backcrossed on the C57BL/6J wild-type background (for more than 15 generations). During experiments, mice were individually housed in custom Plexiglas recording chambers at a constant temperature (23 ± 1°C), humidity (40-60%), and 12/12 light/dark cycle. Food (Laboratory Rodent Diet #5001) and water were available *ad libitum*. All experiments were performed in accordance with the guidelines described in the US National institutes of Health Guide for the Care and Use of Laboratory Animals and were approved by Stanford University’s Administrative Panel on Laboratory Animal Care.

### Virus preparation

Cre-dependent recombinant AAV vectors (pAAV-Ef1α-DIO-GCaMP6s, pAAV-Ef1α-DIO-GFP, and pAAV-Ef1α-DIO-GCaMP6f) were serotyped with AAV-DJ coat proteins and packaged by the Gene Vector and Virus Core at Stanford University. The final viral concentration was 1-1.5 x 10^13^ genome copies (gc) ml^-1^. Aliquots of virus were stored at -80°C until stereotaxic injection.

### Surgery

At the start of surgical procedures, mice were anesthetized with ketamine and xylazine (100 and 20 mg kg^-1^, respectively; IP) and placed into a stereotaxic frame (David Kopf Instruments, Tujunga, CA, USA). To selectively target VTA-GABAergic neurons, virus was infused into the VTA (AP -3.3 mm; ML = 0.2 mm; DV = -4.6 mm) using a 33-guage internal cannula (Plastics One, Inc., Roanoke, VA) attached to a 10 µl Hamilton syringe, at a rate of 0.1 µl min^-1^ (400 nl total virus). After infusion, the internal cannula was kept in place for at least 8 min and then slowly withdrawn. Following virus injections, a 400 µm diameter monofiberoptic cannula (Doric lenses, Inc. Quebec, Canada) was lowered into the VTA (AP = -3.3, ML = 0.3; DV -4.3 mm) for fiber photometry recordings. Mice were also fitted with two gold-plated miniature screw electrodes (AP = 1.5 mm, ML = -1.5 mm and AP = -3.5 mm, ML = -2.8 mm) and two EMG wire electrodes (316SS/44T, Medwire), previously soldered to a four-pin connector (custom). These electrodes were inserted between the trapezius muscles for EMG measurements. The implant was secured to the skull with C&B Metabond (Parkell) and dental cement. The skin was closed with surgical sutures, and mice were maintained on a heating pad until fully mobile. Mice were allowed to recover from surgery for three weeks, and then acclimated to a flexible EEG/EMG connection cable and optical patch cord for ∼10 d within individual recording chambers.

### Fiber photometry recordings and data analysis

Fiber photometry recordings were conducted as previously described (Lerner et al., 2015; Eban-Rothschild et al., 2016). We sinusoidally modulated blue light (30-50 µW) from a 470-nm excitation LED (M470F3, Thorlabs, NJ, USA) at 211 Hz, using a custom Matlab script (MathWorks, Natick, MA, USA) and a multifunction data acquisition device (NI USB-6259, National Instruments, Austin, TX, USA). The blue light was passed through a GFP excitation filter (MD498, Thorlabs), reflected off a dichroic mirror (MD498, Thorlabs), and coupled using a fiber collimation package (F240FC-A, Thorlabs) into a low-fluorescence patch cord (400 µm, 0.48 NA; Doric Lenses) connected to the implanted optic fiber (400 µm) by a zirconia sleeve. GCaMP6 signal was collected through the excitation patch cord, passed through a GFP emission filter (MF525-39, Thorlabs), and focused onto a photodetector (Model 2151, Newport, Irvine, CA, USA) using a lens (LA1540-A, Thorlabs). The signal was then sent to a lock-in amplifier (30-ms time constant, Model SR830, Stanford Research Systems, Sunnyvale, CA, USA) that was synchronized to 211 Hz. Signals from the amplifiers were collected at 1 kHz using a custom Matlab script and a multifunction data acquisition device (National Instruments).

The photometry signal was down-sampled using interpolation to match the EEG sampling rate of 256 Hz. For each experiment, the photometry signal *F* was converted to Δ*F/F* as in Eban-Rothschild et al., 2016. To address a short decay at the beginning of some recording sessions, the Δ*F/F* signal from each session was fitted with a decreasing exponential of the form *a* x *e^bx^*, where *a* > 0 and *b* < 0. We then subtracted the exponential fit from the original Δ*F/F* signal. To detect calcium transients, we generated two filtered Δ*F/F* signals, one low-pass filtered at 0-0.4 Hz, and the other low-pass filtered at 0-40 Hz. We calculated a trace of the derivative of their squared difference and identified candidate transient times by thresholding this signal at mean +1.5 standard deviations. We further required that the Δ*F/F* of candidate transient times be higher than the mean +1.5 standard deviations of the trace. The exact peak time of each transient was identified by taking the maximal value of the thresholded signal within each transient. For each session and for each arousal state, we calculated the mean transient rate (calculated as the number of transients per second) and mean transient amplitude (calculated as the mean Δ*F/F at transient peak times)*.

For sleep-wake analyses of Vgat-cre mice, we recorded data during 7-8 sessions per mouse (n=5), each 30-40 minute long, during the light phase. For sleep-wake analyses of Th-cre mice, we recorded data during 8 sessions per mouse (n=4), each 10-45-min long during the light phase. We used only sessions during which mice showed all three arousal states (wake, NREM, and REM sleep). For each session we calculated the mean Δ*F/F* during all times of wake, NREM sleep, and REM sleep. For state transition analyses, we identified all points of state transition and aligned Δ*F/F* around these times. Mean Δ*F/F* and transient rate of the Th-Cre mice analyzed in this paper were previously reported in Eban-Rothschild et al. 2016.

### EEG/EMG recording and analysis

EEG/EMG signals were amplified (Grass Technologies) and digitized at 256 Hz using Vital Recorder software (Kissei Comtec America). Using analysis software (SleepSign for Animal, Kissei Comtec America) we filtered the signal (EEG: 0.3-80 Hz, EMG: 25-50 Hz) and spectrally analyzed it by fast Fourier transformation to determine a single arousal state (wake, NREM sleep or REM sleep) for each 4-s epoch of data. Wakefulness was defined as desynchronized low-amplitude EEG and heightened tonic EMG activity with phasic bursts. NREM sleep was defined as synchronized, high-amplitude, low-frequency EEG and substantially reduced EMG activity compared with wakefulness, with no phasic bursts. REM sleep was defined as having a pronounced theta rhythm (5-9 Hz) with no EMG activity. The first epoch during which a pronounced theta rhythm was detected, together with reduced EEG amplitude and EMG integral, was defined as the epoch of NREM to REM transition. For all further EEG data analyses, raw (not filtered) EEG/EMG data was exported from the sleep analysis software (SleepSign for Animal) and imported into Matlab using a custom script. To estimate EEG power in a specific band we filtered the EEG in that band and took the absolute value of the Hilbert transform of the filtered trace.

### Salient stimuli presentation

We gently introduced salient stimuli to the home cage of test mice that were individually housed and connected to fiber optic and EEG/EMG recording cables. In each experimental session, each test mouse was exposed to one stimulus (n = 4-5 per stimulus). Different stimuli were presented several days apart. Salient stimuli were: (1) 8 to 12 weeks-old female C57BL/6J mice (Jackson Laboratory), (2) ex-breeder CD1 male mice (>4 month old), (3) 4-weeks-old male C57BL/6J mice, (4) high-fat chow (D12451, Research Diets, Inc.); and (5) rat bedding. We provided test mice with a small piece (0.3 g) of high-fat chow during the two days preceding the high-fat chow experiment. We recorded fiber photometry and EEG/EMG signal prior to and following the introduction of the salient stimuli.

### Immunohistochemistry

Following completion of experiments, mice were deeply anesthetized with ketamine and xylazine (100 and 20 mg kg−1, respectively, i.p.) and transcardially perfused with 1× PBS followed by 4% paraformaldehyde (PFA) in PBS. Brains were removed and post-fixed overnight in 4% PFA. Brains were then cryoprotected in 30% sucrose for minimum 48 h at 4 °C, and then sectioned (30 µm) on a cryostat (Leica Microsystems), collected in PBS containing 0.1% NaN_3_ and stored at 4 °C.

Sections underwent immunohistochemical staining as described below. Primary antibodies were used at 1:1,000 dilution: chicken anti-tyrosine hydroxylase (#TYH, Aves Labs, Inc.) and rabbit anti-GABA (#20094, IMMUNOSTAR). Immunofluorescent secondary antibodies were used at 1:500 dilution: donkey anti-chicken Alexa Fluor 594 (#703-585-155, Jackson ImmunoResearch Laboratories, Inc.) for GCaMP6 and TH co-localization, and donkey anti-rabbit Alexa Fluor 594 (#711-585-152, Jackson ImmunoResearch Laboratories, Inc.) for GCaMP6 and GABA co-localization.

For immunofluorescence staining, brain sections were washed in PBS for 5 min and then incubated in a blocking solution composed of PBS with 0.3% Triton X-100 (PBST) containing 4% bovine serum albumin for 1 h. Sections were then incubated for ∼16 h in block solution containing primary antibodies. After three 5-min washes in PBS, sections were incubated in blocking solution for 1 h and then incubated for 2 h in blocking solution containing secondary antibodies. Sections were washed three times for 5 min each in PBS, mounted onto gelatin-coated microscope slides (FD Neurotechnologies, #PO101) and coverslipped with Fluoroshield containing DAPI Mounting Media (Sigma, #F6057).

### Microscopy and image analysis

Images were collected on a Zeiss LSM 710 confocal fluorescent microscope using reflected light. Digital images were minimally processed using ImageJ (NIH) to enhance brightness and contrast for optimal representation.

### Statistical analyses

Sample sizes were chosen based upon previous publications for the study of the sleep-wake circuitry. We used two-way RM ANOVAs for analyses of calcium signal and transient rate (Figs 1E,G and 5f). We used two-tailed paired t-test’s for data presented in Figures 1F,H and Supplementary Figure 1A. We used correlation coefficient to determine the correlation between fluorescence and power in specific frequency bands (Figs. 3-5 and S2-4). To correct for the multiple comparisons in the analyses of correlation between fluorescence and power in specific frequency bands we used a Bonferroni correction, i.e. multiplied the obtained p-values by the number of comparisons (Wake, x2 (theta and gamma comparisons); NREM, x3 (delta, theta and sigma comparisons); REM, x2 (theta and gamma comparisons)). We also corrected for multiple comparisons by multiplying the obtained p-values by the number of comparisons (3) in the data presented in Fig. S1C. We analyzed data using Prism 6.0 (GraphPad Software) and Matlab R2016b software. Data are presented as mean ± s.e.m. unless otherwise indicated. For the preparation of the figures, we used Matlab R2016b and Prism 6.0 and then exported the data into Adobe Illustrator CS6 (Adobe Systems).

**Figure 1:**
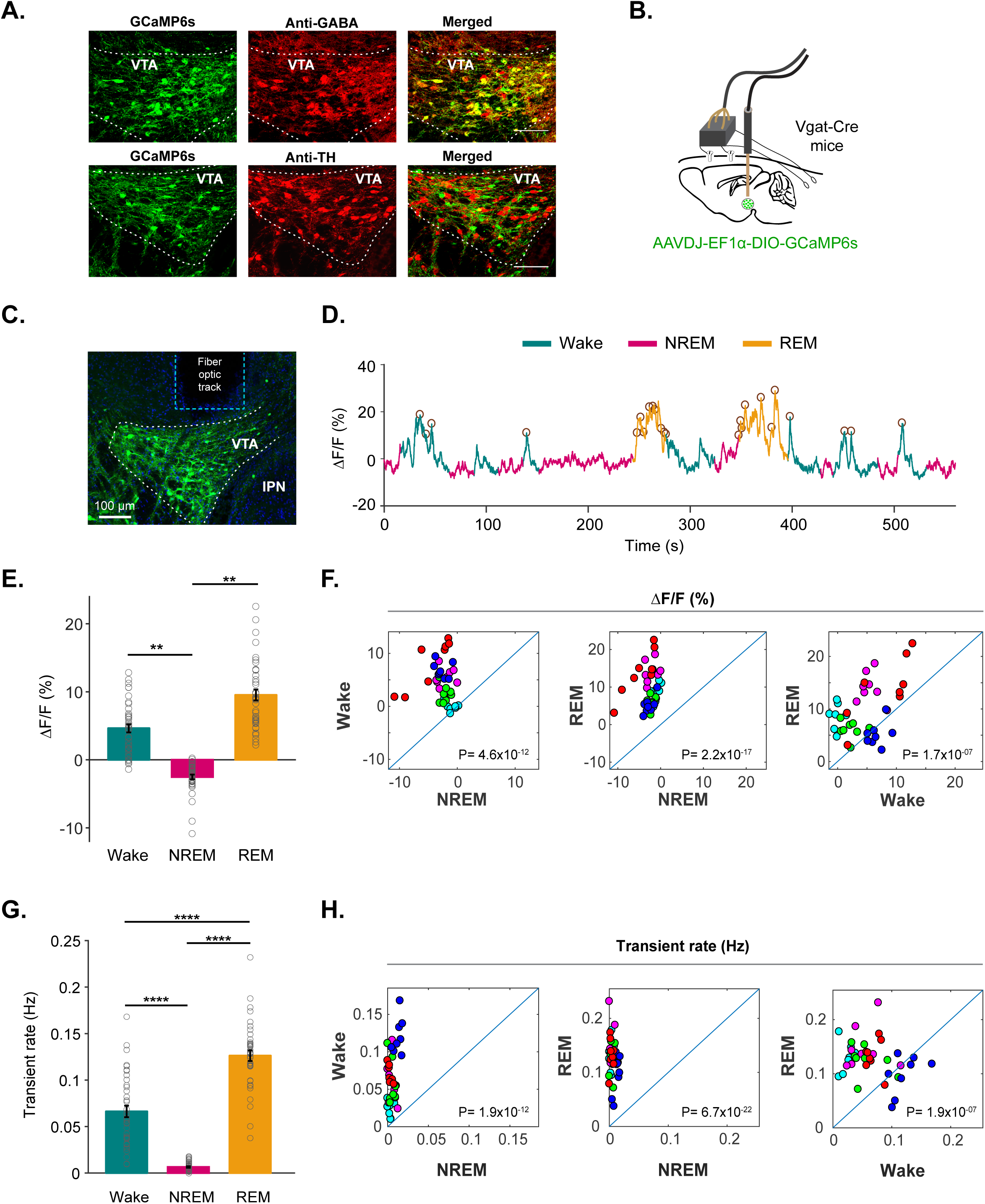
VTA-GABAergic neurons show arousal-state dependent alterations in population activity. (**A**) Confocal microscopic images of AAV-DJ-EF1α-DIO-GCaMP6s expression in the VTA co-immunostained with anti-GABA (top) or anti-TH (bottom) antibodies, demonstrating selective calcium indicator expression in VTA-GABAergic neurons, but not VTA-dopaminergic neurons. (**B**) Schematic for simultaneous recordings of VTA-GABA-GCaMP6s and EEG/EMG signals in *Vgat*-Cre mice. (**C**) Representative image of GCaMP6s-expressing neurons in the VTA and the fiber optic tract for recording. (**D**) Representative fluorescence trace color-coded by arousal state (wake, NREM and REM sleep). Identified transients are denoted by open circles. (**E**) Mean ± s.e.m. fluorescence during wake, NREM and REM sleep (two-way repeated-measures (RM) ANOVA between ‘Mice’ and ‘Arousal state’. Mice, *F_7,31_* = 0.006, *P* >0.99. Arousal state, *F_2,62_* = 8.075, *P* =0.0008, followed by Tukey’s *post hoc* tests, wake vs. NREM *P* >0.99, wake vs. REM *P* =0.0027, NREM vs. REM *P* =0.0027). (**F**) Scatter plots comparing mean %ΔF/F values for pairs of arousal states (paired samples t-test’s, Wake vs. NREM, t(38)= -9.8881, *P*= 4.6×10^-12^; REM vs. NREM, t(38)= -14.8166, *P*= 2.2×10^-17^; Wake vs. REM, t(38)= -6.387, *P*= 1.68×10^-7^). Each dot represents a single trial. The different colors represent different mice. (**G**) Transient rate (Hz) for wake, NREM and REM sleep (two-way repeated-measures (RM) ANOVA between ‘Mice’ and ‘Arousal state’. Mice, *F_7,31_* = 0.7776, *P* =0.61. Arousal state, *F_2,62_*= 117.8, *P* < 1×10^-15^, followed by Tukey’s *post hoc* tests, wake vs. NREM = 4.39×10^-10^, wake vs. REM *P* = 3.99×10^-10^, NREM vs. REM *P* = 1.89×10^-11^). (H) Scatter plots comparing mean transient rate values among each pair of arousal state (paired samples t-test’s, Wake vs. NREM, t(38)= -10.2078, *P*= 1.9×10^-12^; REM vs. NREM, t(38)= -20.1677, *P*= 6.7×10^-22^; Wake vs. REM, t(38)= -6.3469, *P*= 1.9×10^-7^). Each dot represents a single trial. The different colors represent different mice.

## Results

### VTA-GABAergic neuronal activity is modulated across arousal state

To assess population activity of VTA-GABAergic neurons across spontaneous sleep-wake states we used calcium-based fiber photometry recordings (**Fig. 1A-C**). We injected a Cre-dependent adeno-associated virus (AAV) encoding the fluorescent calcium indicator GCaMP6s (pAAV-DJ-EF1α-DIO-GCamP6s) into the VTA of Vgat-IRES-Cre knock-in mice and implanted a fiber optic probe above the VTA for subsequent delivery of excitation light and collection of fluorescent emission and with EEG-EMG electrodes for simultaneous sleep-wake recordings. Fluorescent immunostaining for GABA and tyrosine hydroxylase (TH) in GCaMP6-expressing slices demonstrated specific co-expression of GCaMP6 in GABAergic, but not dopaminergic VTA neurons (**Fig. 1A**). We found robust alterations in both fluorescence signal (%ΔF/F) and calcium transient rate (Hz) in VTA GABAergic neurons over spontaneous sleep/wake states (**Fig. 1D-H**). During NREM sleep, VTA-GABAergic neurons showed the lowest fluorescence and calcium transient rates compared to both wakefulness and REM sleep (**Fig. 1D-H**). In contrast, VTA GABAergic neurons showed the largest and most frequent calcium transients during REM sleep compared to both wakefulness and NREM sleep (**Fig. 1G-H** and **Fig. S1A**). These arousal-state-dependent alterations in activity were not found in control experiments (**Fig. S2A-B**).

We proceeded to determine whether activity in VTA-GABAergic neurons is altered around arousal-state transitions. VTA-GABAergic neurons showed a strong decrease in population activity at wake to NREM transitions (**Fig. 2A-B**), while EEG delta power was strongly increased (**Fig. 2C**). These alterations were not found in control experiments (**Fig. S2C**). NREM sleep to wakefulness transitions were accompanied by a large increase in population activity and a decrease in EEG delta power (**Fig. 2A-C**), even for short (<12 sec) wake episodes (**Fig. S1B**). During NREM to REM sleep transitions, the Δ*F/F* signal showed a gradual ramping 10-20 seconds prior to the vigilance state transition, subsequently peaking ∼20 seconds following the transition (**Fig. 2A,B**). Following REM sleep to wake transitions, a reduction in VTA-GABAergic calcium fluorescence was observed (**Fig. 2A,B**). Together, these findings demonstrate that VTA-GABAergic neurons exhibit strong alterations in population activity across arousal states.

**Figure 2:**
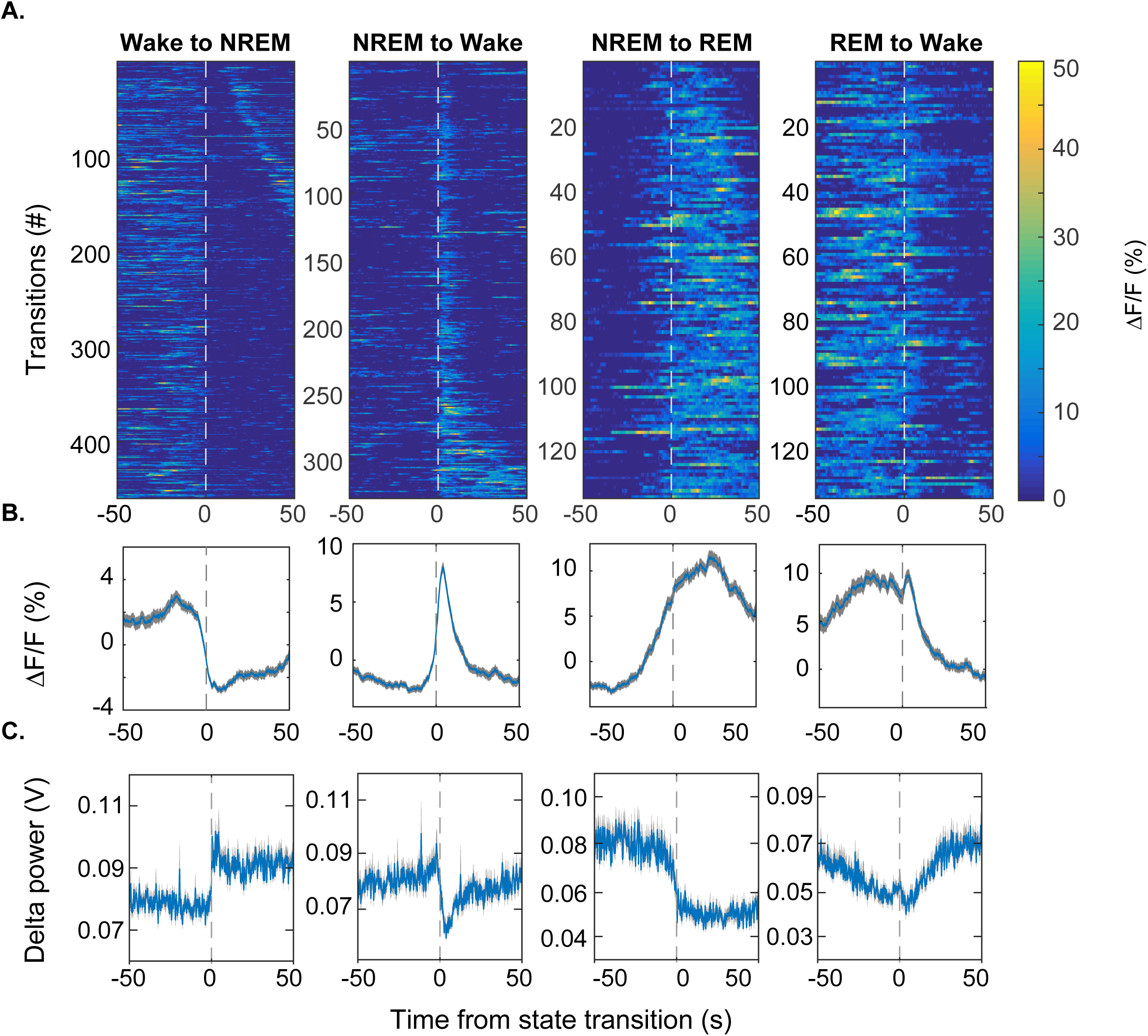
VTA-GABAergic population dynamics around arousal state transitions. (**A**) Heatmaps of fluorescence intensity aligned to arousal state transitions (wake to NREM, *n* = 459; NREM to wake, *n* = 329; NREM to REM, *n* = 135; REM to wake, *n* = 135). Trials are sorted by the duration of the second state (shortest, top; longest, bottom). (**B**) Average responses from all the transitions expressed as mean (blue trace) ± s.e.m. (gray shading). (**C**) EEG delta (1-4.5 Hz) power from all the transitions.

### VTA-GABAergic neuronal activity correlates with specific cortical oscillatory patterns

To investigate whether VTA-GABAergic neural activity is correlated with specific components of the EEG during different vigilance states, we examined correlations between population activity and power within specific EEG bands during 1-sec epochs of wakefulness, NREM, and REM sleep. During wakefulness, we found a negative correlation between VTA-GABAergic population activity and EEG theta (5-9 Hz) power, and a positive correlation of population activity and EEG gamma (30-80 Hz) power (**Fig. 3A-D**). The positive gamma correlation was evident across all test mice (**Fig. S3**), and was absent in control experiments (**Fig. S2D**). In contrast, population activity of VTA-dopaminergic neurons was not correlated with the EEG gamma band, and only weakly correlated with theta (**Fig. 3E-H**). These data suggest that during wakefulness, activity of VTA-GABAergic, but not dopaminergic, neurons is related to fast cortical oscillations.

**Figure 3:**
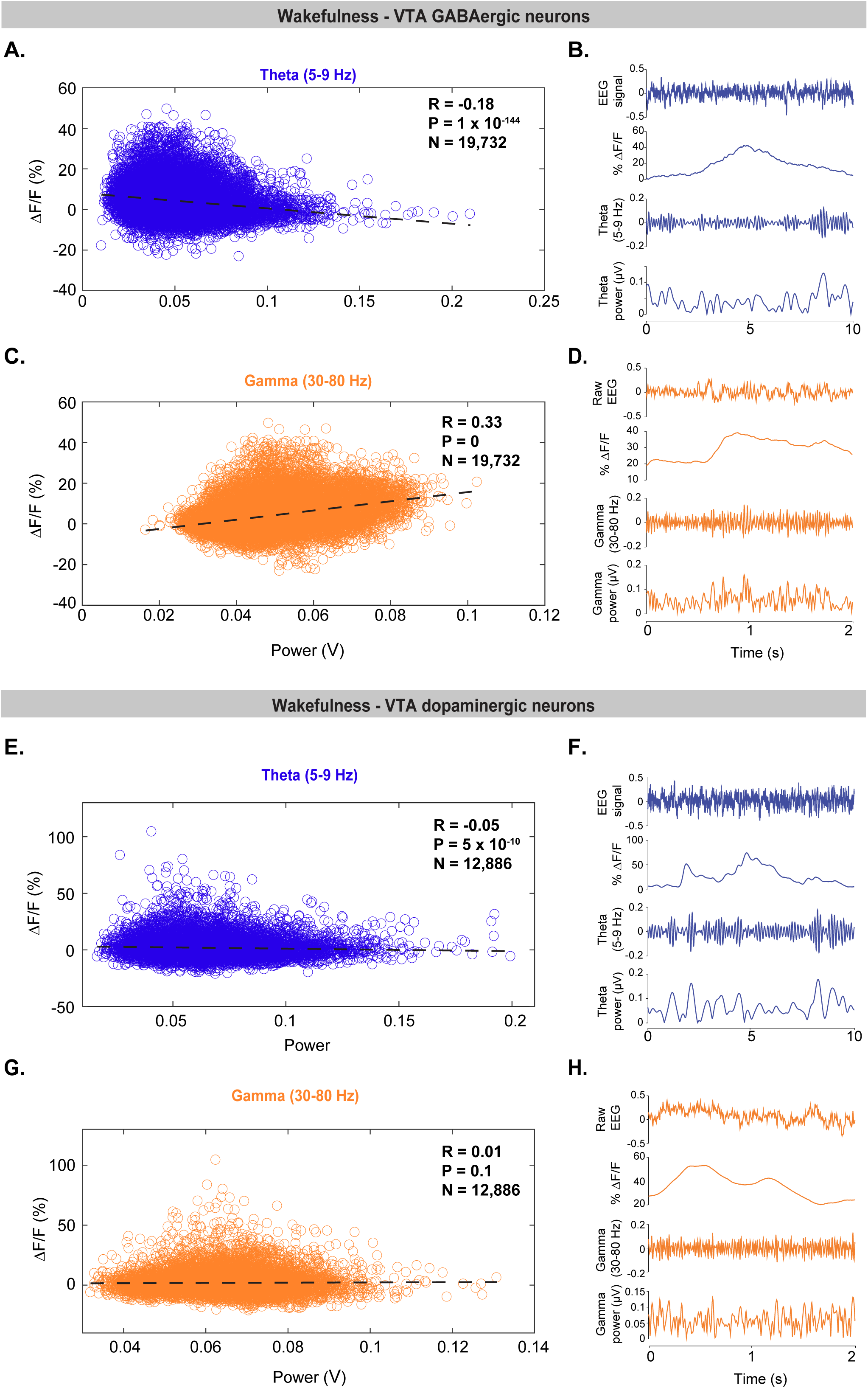
Population activity of VTA-GABAergic, but not dopaminergic, neurons correlates with EEG gamma power during wakefulness. (**A,C,E,G**) Correlations between EEG spectral components and population activity during spontaneous wakefulness. Each dot represents a 1-second epoch. Dashed line is the best linear fit. R is correlation coefficient. (**B,D,F,H**) Representative raw EEG trace, fluorescence trace, filtered EEG and power in a specific EEG band for a 2-10 seconds wake epoch. (**A**) VTA-GABAergic population activity plotted against EEG theta (5-9 Hz) power. (**B**) Representative traces for theta band and VTA-GABAergic population activity. (**C**) VTA-GABAergic population activity plotted against EEG gamma (30-80 Hz) power. (**D**) Representative traces for gamma-band and VTA-GABAergic population activity. (**E**) VTA-dopaminergic population activity plotted against EEG theta power. (**F**) Representative traces for theta band and VTA-dopaminergic population activity. (**G**) VTA-dopaminergic population activity plotted against EEG gamma power. (**H**) Representative traces for gamma-band and VTA-dopaminergic population activity.

During NREM sleep, we observed a negative correlation between VTA-GABAergic population activity and EEG delta (1-4.5 Hz), theta and sigma (10-14 Hz) power bands (**Fig. 4**). Moreover, power in these frequency bands was significantly lower during high Δ*F/F* times as compared to low Δ*F/F* times (**Fig. S1C**). Similar to VTA-GABAergic neurons, VTA-dopaminergic neurons showed a negative correlation between population activity and EEG delta, theta, and sigma power bands during NREM sleep (**Fig. S4A-C**). Together our findings suggest that within NREM sleep, high activity periods in both VTA-GABAergic and VTA-dopaminergic neurons are associated with a lightening of NREM sleep.

**Figure 4:**
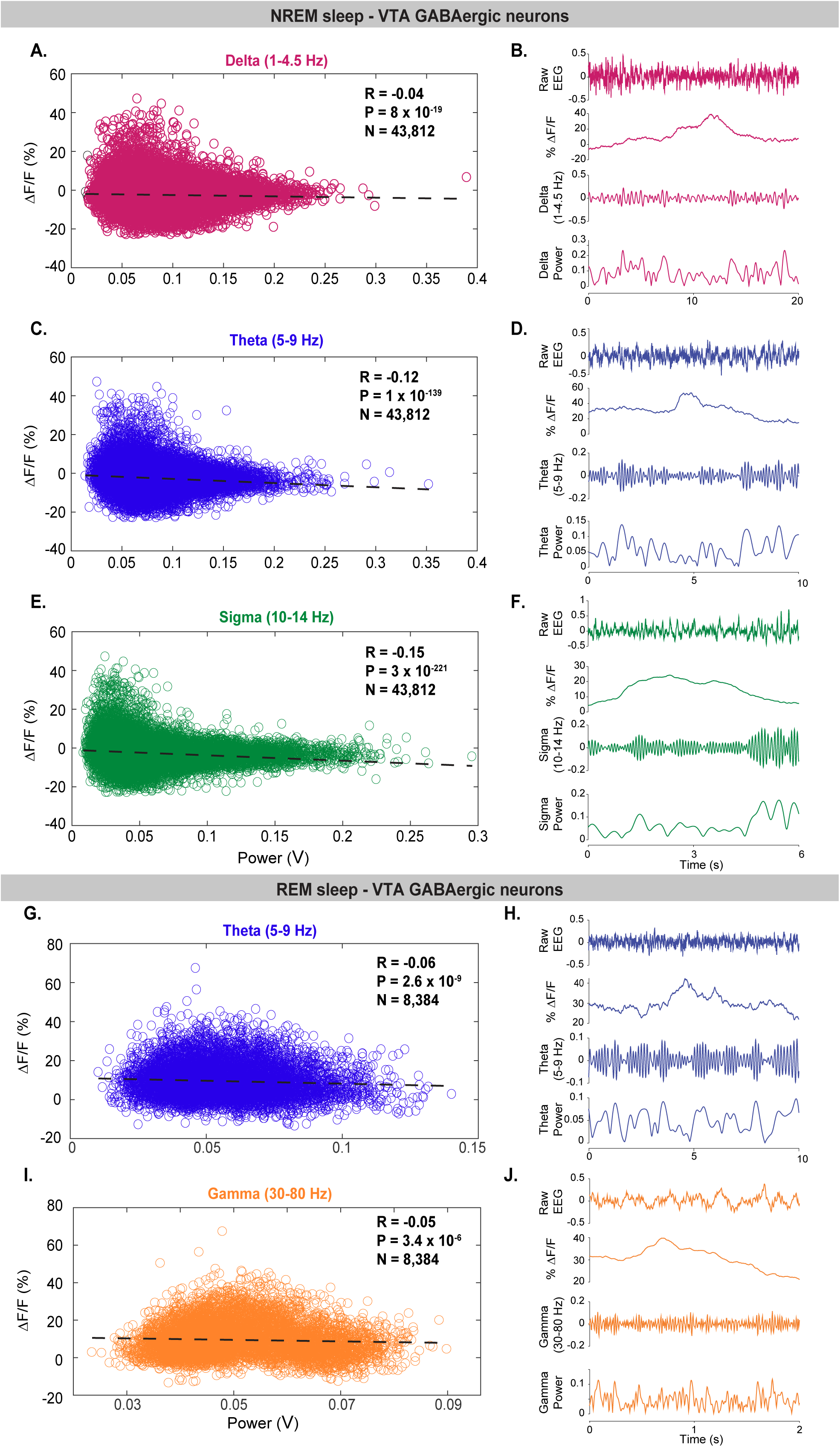
VTA-GABAergic neural activity is negatively correlated with low-frequency EEG oscillations during NREM sleep. (**A-F**) Correlations between EEG spectral components and VTA-GABAergic neural activity during spontaneous NREM sleep. (**A,B**) Delta power correlations and representative data during NREM sleep. (**C,D**) Theta power correlations and representative data during NREM sleep. (**E,F**) Sigma power correlations and representative data during NREM sleep. (**G-J**) Correlations between EEG theta and gamma frequency bands and VTA-GABAergic neural activity during spontaneous REM sleep. (**G,H**) Theta power correlations and representative data during REM sleep. (**I,J**) Gamma power correlations and representative data during REM sleep. Each dot represents a 1-second epoch. Dashed line is the best linear fit. R is correlation coefficient.

Within REM sleep, we found a significant, yet weak, negative correlation between VTA-GABAergic population activity and both EEG theta and gamma power bands (**Fig. 4G-J**). In contrast, VTA-dopaminergic neurons showed an opposite, yet weak, trend for EEG theta power and no correlation for EEG gamma power (**Fig. S4D-F**). Taken together, these findings demonstrate that activity in VTA-GABAergic neurons is strongly associated with both behavioral and electrocortical arousal.

### VTA-GABAergic neurons exhibit biased responses to social stimuli

To further explore the relationship between activity in VTA-GABAergic neurons and arousal, we presented mice with various salient stimuli, of both positive and negative valences, while simultaneously recording population activity from VTA-GABAergic neurons and EEG/EMG signals. Following the placement of salient stimuli inside the home cage of test mice, VTA-GABAergic neurons showed a fast increase in population activity (**Fig. 5A-E**). Social stimuli (female mouse, juvenile mouse and an aggressive mouse) induced an increase in activity that remained elevated for more than 120 seconds, irrespective of valence (**Fig. 5A-C**). Non-social salient stimuli, associated with both positive (high fat chow) and negative (rat bedding) valences, induced much weaker activation of VTA-GABAergic neurons (**Fig. 5D-F**). As during spontaneous wakefulness, population activity of VTA-GABAergic neurons showed a positive correlation with EEG gamma power during the presentation of most salient stimuli (**Fig. 5G-K**). Surprisingly, no correlation between population activity of VTA-GABAergic neurons and EEG gamma power was found during trials in which male mice were presented (both young and older/aggressive males) (**Fig. 5G-K**). Together, these findings further support the premise that VTA-GABAergic neurons activity is tightly linked to behavioral and electrocortical arousal.

**Figure 5:**
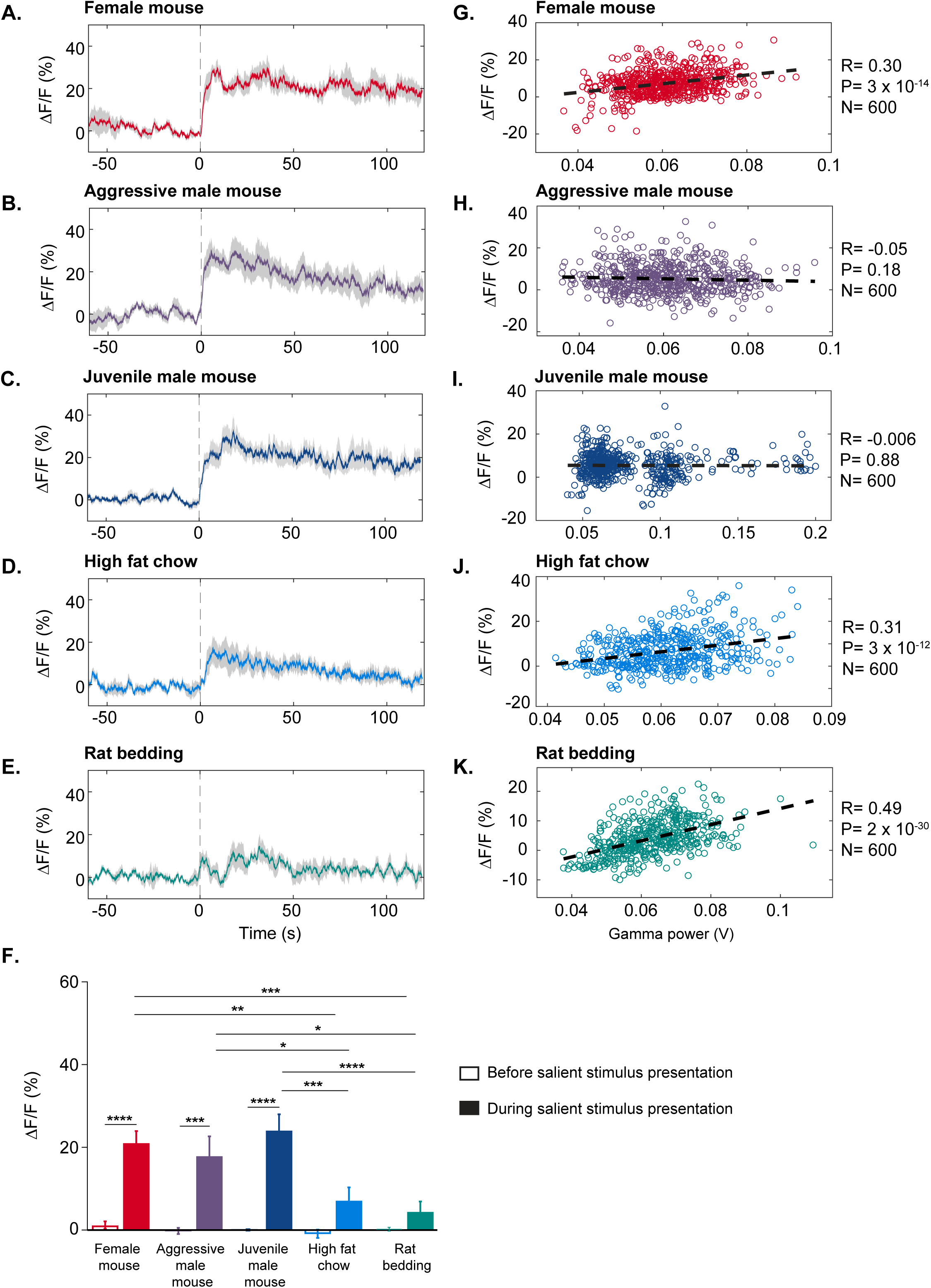
VTA-GABAergic neurons are strongly activated by social salient stimuli. (**A-E**) Population activity of VTA-GABAergic neurons prior and following the placement of (**A**) a female mouse, (**B**) an aggressive (CD1) male mouse, (**C**) a juvenile male mouse, (**D**) high fat rodent chow, and (**E**) rat bedding, inside the home cage of test mice. Presented are 60 seconds prior to and 120 seconds following stimulus presentation. (**F**) Summary data showing the relative change in %ΔF/F for each stimulus prior to and after presentations (two-way ANOVA: main effect of time F_1,19_ = 95.92, p < 0.0001; main effect of stimuli F_4,19_ = 5.324, p = 0.0048; and the interaction F_4,19_ = 5.759, p = 0.0033). Sidak’s multiple comparisons post-hoc tests reveal stimulus-specific responses in the %ΔF/F signal (no differences were observed during baseline prior to stimulus presentation; after stimulus presentation, female mouse vs. aggressive mouse t = 0.8874, p > 0.05, female mouse vs. juvenile male mouse t = 0.8488, p > 0.05, female mouse vs. high fat diet t = 3.901, p = 0.0038, female mouse vs. rat (predator) bedding t = 4.391, p = 0.0009, aggressive mouse vs. juvenile mouse t = 1.736, p > 0.05, aggressive mouse vs. high fat diet t = 3.014, p = 0.0448, aggressive mouse vs. rat bedding t = 3.555, p = 0.0103, juvenile mouse vs. high fat diet t = 4.75, p = 0.0003, juvenile mouse vs. rat bedding t = 5.192, p < 0.0001, high fat diet vs. rat bedding t = 0.7133, p > 0.05. Before and after stimulus comparisons further allow for specific effects of each stimulus to be assessed (Sidak’s): female mouse t = 5.983, p < 0.0001, aggressive mouse t = 5.459, p = 0.0001, juvenile mouse t = 7.249, p < 0.0001, high fat diet t = 2.474, p > 0.05, rat bedding t = 1.14, p > 0.05. Note that before and after stimulus differences are only significant for social stimuli (female and male mice), but not for nonsocial stimuli (high fat chow and rat bedding). **(G-K)** Correlations between EEG gamma power and VTA-GABAergic neural activity following the presentation of salient stimuli. Each dot represents a 1-second epoch. Dashed line is the best linear fit. R is correlation coefficient.

## Discussion

Here, we investigated the relationships between electrocortical and behavioral arousal and population activity of subsets of neurons in the VTA. We observed that activity in VTA-GABAergic neurons varies significantly across arousal states. VTA-GABAergic neurons are strongly activated during wakefulness and REM sleep and show low population activity during NREM sleep. Moreover, during NREM sleep, population activity of VTA-GABAergic neurons negatively correlates with NREM sleep intensity, i.e. delta, theta and sigma power bands. In contrast, during wakefulness population activity of VTA-GABAergic neurons is positively correlated with EEG gamma power. We also found that VTA-GABAergic neurons are strongly activated by different salient stimuli, specifically social stimuli of both positive and negative valence. Activity of VTA-dopaminergic neurons was found to be similarly negatively correlated with NREM sleep intensity but, notably, we did not observe a significant correlation between population activity and gamma power during wakefulness. Together, our findings demonstrate that the VTA, which is mainly studied in regard to reward, motivation and learning, also shows strong modulation by arousal state.

Both VTA dopaminergic and GABAergic neurons are strongly activated by rewards and reward-predictive cues, yet VTA-GABAergic neurons show sustained excitation following a reward-predictive cue while activity in VTA-dopaminergic neurons is suppressed in the delay period between cue and reward delivery (Cohen et al., 2012). It has been suggested that sustained activation of VTA-GABAergic neurons during the delay period accounts for the suppression of VTA-dopaminergic neural activity and possibly the computation of reward prediction error (Schultz, 1998; Cohen et al., 2012). VTA-GABAergic neurons inhibit VTA-dopaminergic neurons both *in vitro* (Dobi et al., 2010; van Zessen et al., 2012) and in anesthetized rats (Tan et al., 2012). It has been proposed that VTA-GABAergic neurons do not exert a tonic inhibition on VTA-dopaminergic neurons, but rather sculpt their activity pattern in favor of fast oscillations (Brown and McKenna, 2015). In the context of sleep/wake regulation, we have previously demonstrated that VTA-dopaminergic neurons increase their activity prior to NREM sleep-to-wake transitions (Eban-Rothschild et al., 2016), while here we report that VTA-GABAergic neurons increase their activity in conjunction with these transitions (Fig. 2). It is thus unlikely that VTA-GABAergic neurons drive NREM sleep-to-wake transitions. Rather, VTA-GABAergic neurons may suppress activity in VTA-dopaminergic neurons following a sleep-to-wake transition–to tightly control arousal.

Indeed, lesioning or chemogenetic inhibition of VTA-GABAergic neurons produces persistent wakefulness (Takata et al., 2018; Yu et al., 2019b). VTA-dopaminergic neural activation potently promotes arousal and maintains long-term wakefulness (Eban-Rothschild et al., 2016; Taylor et al., 2016; Oishi et al., 2017), and the increased activity in VTA-dopaminergic neurons suppresses the motivation to prepare for sleep and engage in sleep-preparatory behaviors (Eban-Rothschild et al., 2016). It is reasonable to assume that activity in VTA-dopaminergic neurons is tightly regulated to suppress wakefulness when sleep is in need and prevent a state of hyperarousal (Yu et al., 2019a)–as may be the case in mania or drug-induced arousal. Cocaine, for example, reduces activity in VTA-GABAergic neurons, which results in disinhibition of VTA-dopaminergic neurons and the elevation of extracellular dopamine concentrations (Bocklisch et al., 2013). Similarly, optogenetic inhibition of VTA-GABAergic neurons results in disinhibition of VTA-dopaminergic neurons and an increase in VTA-dopaminergic neuronal activation (Bocklisch et al., 2013). Together, our findings provide a mechanistic framework for the participation of dopaminergic and GABAergic VTA neurons in the regulation of arousal. A better understanding of VTA microcircuitry in the context of arousal regulation holds great promise to improve the capacity to treat sleep dysregulation in pathological circumstances, such as drug addiction.

It is important to mention that in the current study we did not distinguish between subpopulations of GABAergic neurons. For example, some VTA-GABAergic neurons express corticotropin-releasing factor (CRF)-binding protein (Wang and Morales, 2008), dopamine receptor D2, or µ-opioid receptors (Margolis et al., 2012). In addition, some VTA-GABAergic interneurons form local connections within the VTA while others send long-range projections (Morales and Margolis, 2017). It would be of interest for future studies to determine whether these disparate populations differentially influence arousal. Recently, Yu and colleagues made progress in this regard. They demonstrated that VTA-GABAergic neurons project to the lateral hypothalamus, where they inhibit wakefulness and drive NREM sleep (Yu et al., 2019b). When VTA-GABAergic neural activity was monitored using fiber photometry, they observed high calcium fluorescence during wakefulness and REM sleep, similar to our findings. In contrast, Chowdhury and colleagues recently observed that GAD67+ GABAergic neurons in the VTA are mostly active during NREM sleep (Chowdhury et al., 2019). The discrepancy between the different studies may be due to the use of a different Cre-line to target GABAergic neurons in the VTA (Vgat-cre vs. GAD67-cre), or due to differences in data interpretations. Although Chowdhury and colleagues reported higher population activity in VTA GAD67+ neurons during NREM sleep compared to wakefulness and REM sleep, examination of their data suggests an increase in population activity immediately following sleep-to-wake transitions, and higher transient rates during wakefulness and REM sleep compared to NREM sleep (Chowdhury et al., 2019). Therefore, additional work needs to be done to tease apart how specific subsets of VTA-GABAergic neurons contribute to arousal state regulation.

How are VTA-GABAergic neurons linked to other sleep/wake regulatory centers? VTA-GABAergic neurons receive excitatory, inhibitory and modulatory inputs from diverse cortical and subcortical areas (Beier et al., 2015; Faget et al., 2016). Putative VTA-GABAergic neurons are strongly activated by different arousal-related neuromodulators including hypocretin, CRF, substance P, and histamine (Korotkova et al., 2002; Korotkova et al., 2006). VTA-GABAergic neurons send widespread projections to various forebrain and brainstem areas including the basal forebrain, lateral hypothalamus, periaqueductal gray and dorsal raphe nuclei, as well as sparser projections to the nucleus accumbens and prefrontal cortex (Carr and Sesack, 2000b; Kirouac et al., 2004; Brown et al., 2012; van Zessen et al., 2012; Taylor et al., 2014). Together these observations demonstrate that VTA-GABAergic neurons are well positioned to regulate local and global neuronal activity for the support of electrocortical and behavioral arousal.

We found that VTA-GABAergic neural activity correlated with specific components of the EEG power spectrum. During NREM sleep, delta, theta and sigma power bands were negatively correlated with VTA-GABAergic activity (Fig. 4). Slow oscillations in the EEG are related to the depth of sleep, correlate with the duration of prior waking, and are a signature of widespread synchronization of cortical and thalamo-cortical networks (Contreras et al., 1996; Cirelli, 2009; Borbely et al., 2016). Sleep spindles, periodic oscillatory patterns in the NREM EEG that overlap with the sigma power band (10-14 Hz), are thought to stabilize sleep and promote memory consolidation (Mednick et al., 2013; Rasch and Born, 2013). Spindles are primarily generated by the thalamic reticular nucleus (RTN), where GABAergic neurons function as a pacemaker, and glutamatergic thalamo-cortical projections mediate the propagation of spindles to the cortex (von Krosigk et al., 1993; Contreras et al., 1996; De Gennaro and Ferrara, 2003). A similar pattern was observed for VTA-dopaminergic neurons, where high activity within this population was negatively correlated with EEG delta, theta and sigma bands during NREM sleep. Negative correlations between VTA neural activity and cortical NREM sleep characteristics suggest that the concerted activity of these neurons may fundamentally oppose slow oscillations during NREM sleep. This may underlie chronic sleep disturbances in people who abuse drugs targeting the dopaminergic system (e.g., cocaine, methamphetamine).

We also found that activity in VTA-GABAergic, but not dopaminergic neurons, is positively correlated with EEG gamma power during both spontaneous and motivated wakefulness (Fig. 3 and 5). A previous study demonstrated that activity in the VTA and prefrontal cortex (PFC) is coupled via a 4 Hz oscillation that synchronizes local gamma oscillations and neuronal firing (Fujisawa and Buzsaki, 2011). Moreover, the authors reported a task-dependent increase in gamma coherence between the PFC and the VTA during a working memory task (Fujisawa and Buzsaki, 2011). The cellular identity in the VTA accounting for this coupling remained elusive (Fujisawa and Buzsaki, 2011). VTA-GABAergic neurons both receive input from and send reciprocal projections to the PFC (Carr and Sesack, 2000b, a). It is possible that the strong correlation between VTA-GABAergic activation and EEG gamma oscillations reflects a top-down cognitive process via transmission of information from the PFC to fast-spiking VTA-GABAergic neurons. Cortical gamma band activity is largely generated by fast-spiking GABAergic parvalbumin positive interneurons (Sohal et al., 2009), and is hypothesized to enhance information processing and attention (Womelsdorf et al., 2007). VTA-GABAergic activity may also be related to cortical processes associated with gamma activity, such as the temporal representation and consolidation of information (Buzsaki and Draguhn, 2004). Interestingly, our findings suggest that not all behaviors are characterized by this gamma coupling, including interactions with same sex conspecifics. Indeed, further studies using projection-specific manipulations could add valuable insights to the functional role of VTA-GABAergic neural activity and their relation to cortical gamma oscillations.

Lastly, we observed strong and enduring calcium responses from VTA-GABAergic neurons during the presentation of social stimuli. Specifically, large increases in neuronal activity were observed in response to a novel female mouse, aggressive male intruder, and a 4-week old juvenile mouse, suggesting that population-level VTA-GABAergic responses to social stimuli are independent of valence. In contrast, we observed significantly smaller responses to both positive (high fat diet) and negative (rat bedding) stimuli that did not involve a social novelty component. It is important to note that we did not control for possible motor aspects of social stimuli. Although Lee et al. demonstrated that the firing rate of VTA-GABAergic neurons was phasically modulated by certain movements, such as head orienting, little variation in firing rate was observed during sustained locomotor activity (Lee et al., 2001). VTA neurons, specifically dopaminergic, are associated with social reward and social novelty (Gunaydin et al., 2014; Hung et al., 2017), yet the precise function of VTA-GABAergic neurons in this regard awaits further investigation.

In summary, we demonstrated that VTA-GABAergic neural activity is strongly modulated by arousal state. As many addictive drugs inhibit VTA-GABAergic neurons, resulting in VTA dopaminergic disinhibition (Hyman et al., 2006; Luscher and Malenka, 2011) a better understanding of VTA microcircuitry in arousal regulation may lead to novel treatments for diseases involving disorders of cortical activation, hyper-arousal, and disrupted sleep.

---

**Supplementary Figure 1: Population activity of VTA-GABAergic neurons across arousal states.** (**A**) Scatter plots of calcium transient amplitudes during each pair of arousal states (paired samples t-test, Wake vs. NREM, t(33)= -0.9043, *P*= 0.37; REM vs. NREM, t(33)= -3.0933, *P*= 0.004; Wake vs. REM, t(38)= -2.5326, *P*= 0.01). Each dot represents the average across a single trial. The different colors represent different mice. (**B**) Mean (blue trace) ± s.e.m. (gray shading) fluorescence from all NREM to wake transitions when wake episodes are shorter than 12 sec. *n* = 90 (**C**) Distribution of Delta, theta and Sigma power for high (green) and low (orange) %ΔF/F values. ‘High’ %ΔF/F values were defined as >= 3 standard deviations above the mean; ‘Low’ %ΔF/F values were defined as < 3 standard deviations above the mean (student’s t-test: Delta, t(43810)=10.9897; Theta, t(43810)=12.2218; Sigma, t(43810)=14.6082).

**Supplementary Figure 2: Fiber-photometry recordings from a control mouse.** We stereotaxically injected the control virus AAV-DJ-EF1α-DIO-GFP into the VTA of a Vgat-Cre mouse and implanted a fiber optic probe (400 µm) and EEG/EMG electrodes. (**A**) Representative fluorescence trace color-coded by arousal state. (**B**) Mean (± s.e.m.) fluorescence during wake, NREM and REM sleep. *N* = 6 trials. (**C**) Mean (blue trace) ± s.e.m. (gray shading) fluorescence from all wake-to-NREM sleep transitions aligned to arousal state transitions. *N* = 44 transitions. (**D**) Correlations between EEG gamma power and population activity during spontaneous wakefulness. Each dot represents a 1-second epoch. Dashed line is the best linear fit. R is correlation coefficient.

**Supplementary Figure 3: Correlation between EEG gamma power and VTA GABAergic population activity during spontaneous wakefulness for individual mice. (A-E)** Data obtained from mice #1-5.

**Supplementary Figure 4: Correlations between EEG spectral components and VTA-dopaminergic neural activity during NREM and REM sleep.** (**A-C**) Correlations between EEG spectral components and VTA-dopaminergic population activity during spontaneous NREM sleep. (**A**) VTA-dopaminergic population activity plotted against EEG delta power. (**B**) VTA-dopaminergic population activity plotted against EEG theta power. (**C**) VTA-dopaminergic population activity plotted against EEG sigma power. (**D-E**) Correlations between EEG theta and gamma frequency bands and VTA-dopaminergic neural activity during spontaneous REM sleep. (**D**) VTA-dopaminergic population activity plotted against EEG theta power. (**E**) VTA-dopaminergic population activity plotted against EEG gamma power. Each dot represents a 1-second epoch. Dashed line is the best linear fit. R is correlation coefficient.

