## Supplementary figures and images for "Arousal-State Dependent Alterations in VTA-GABAergic Neural Activity"

### Supplementary Figures

Supplementary Figure 1

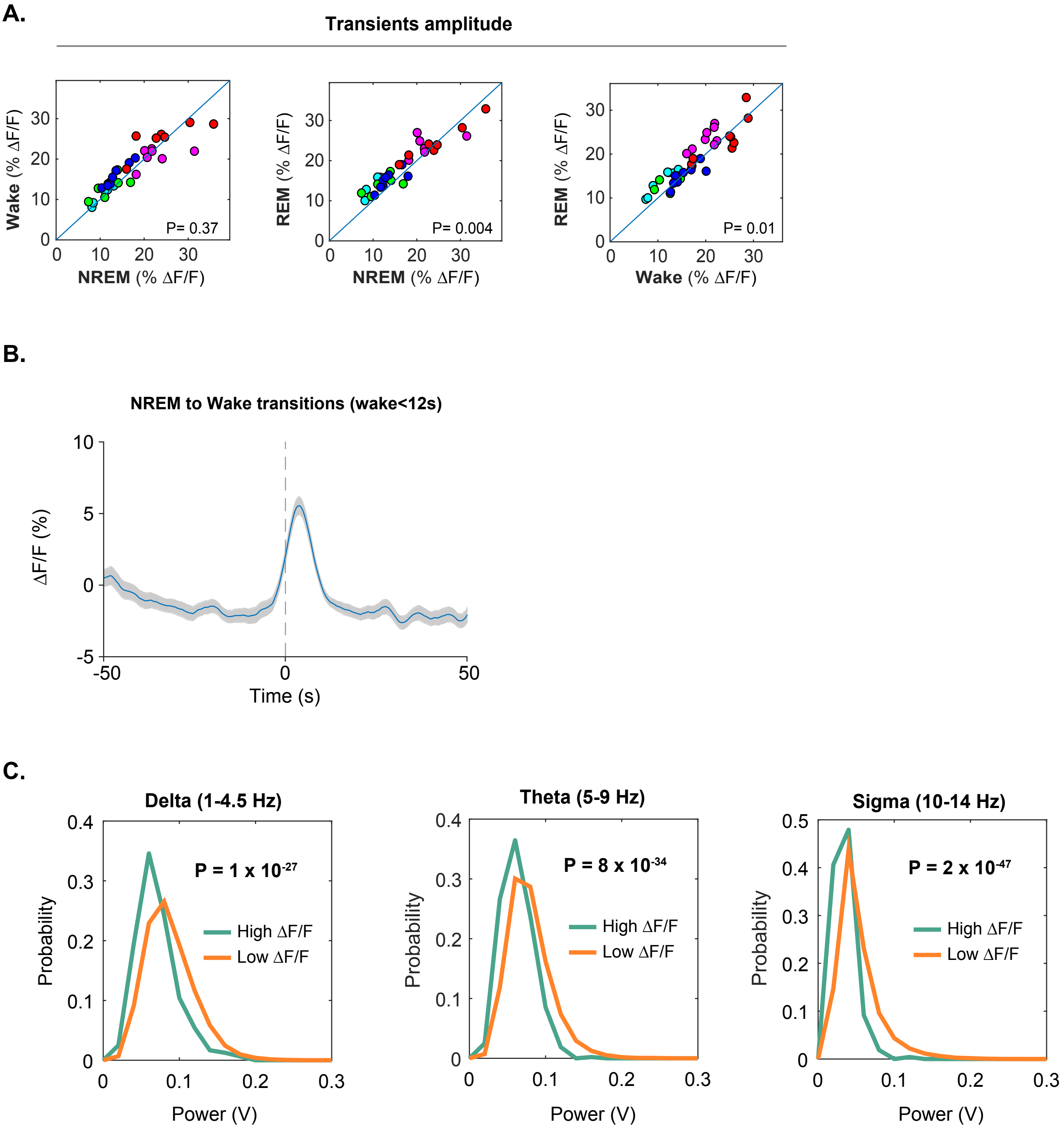

Supplementary Figure 2

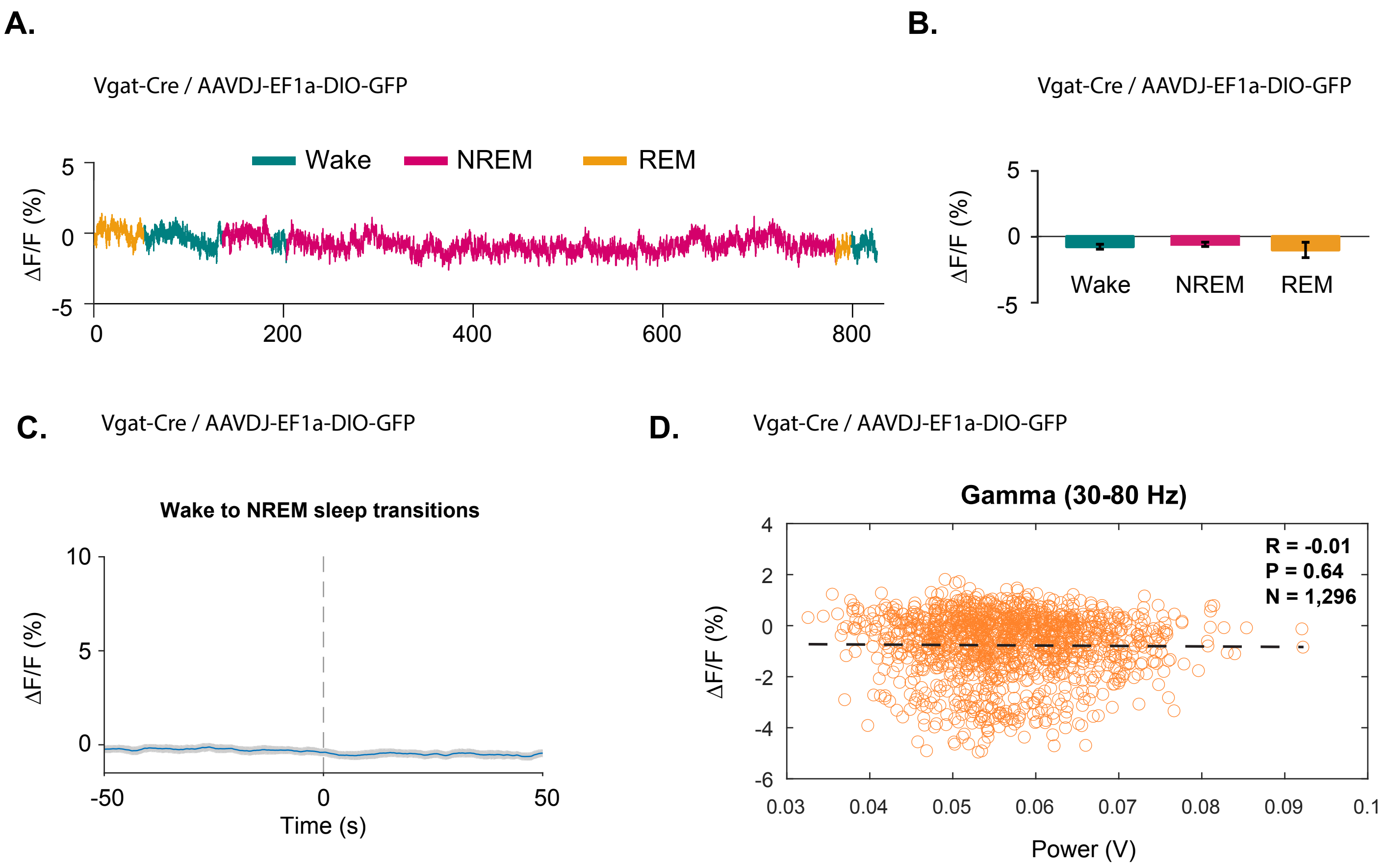

Supplementary Figure 3

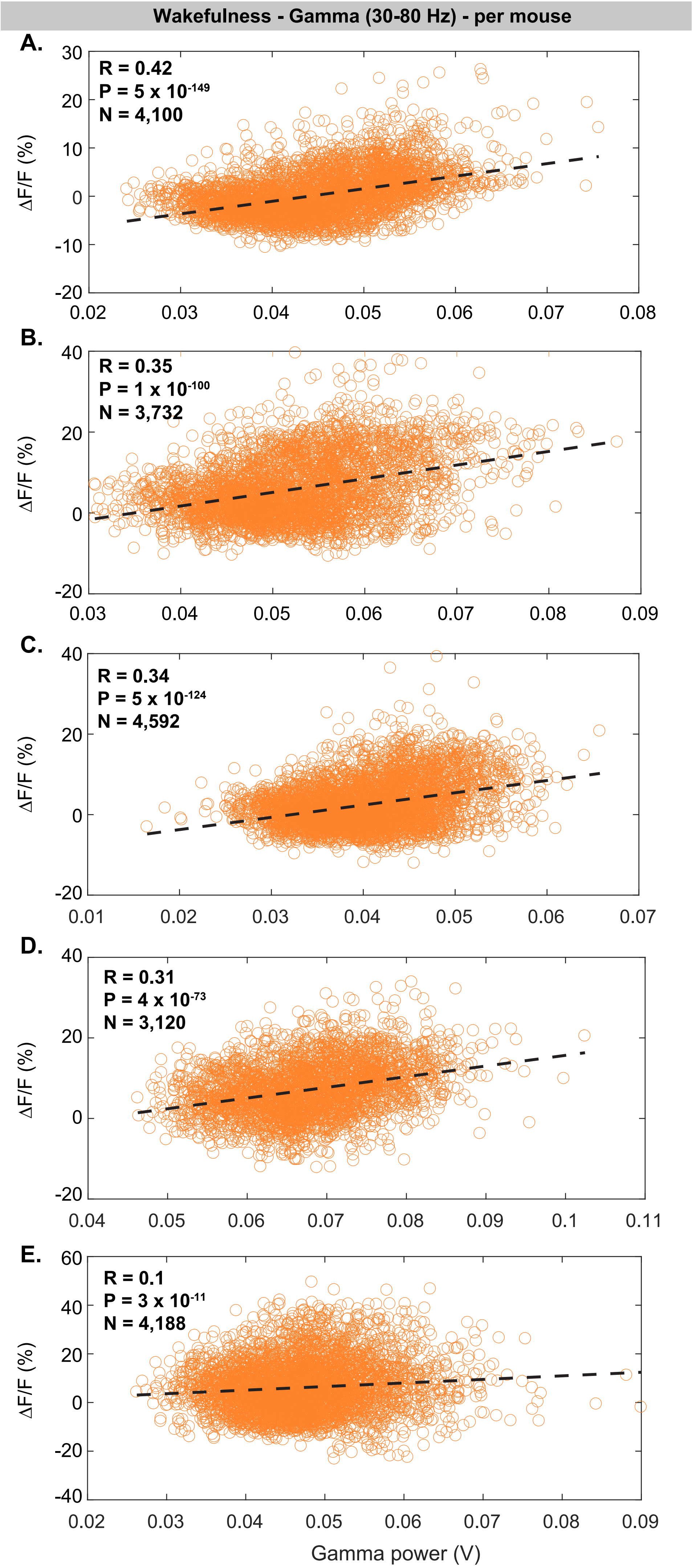

Supplementary Figure 4

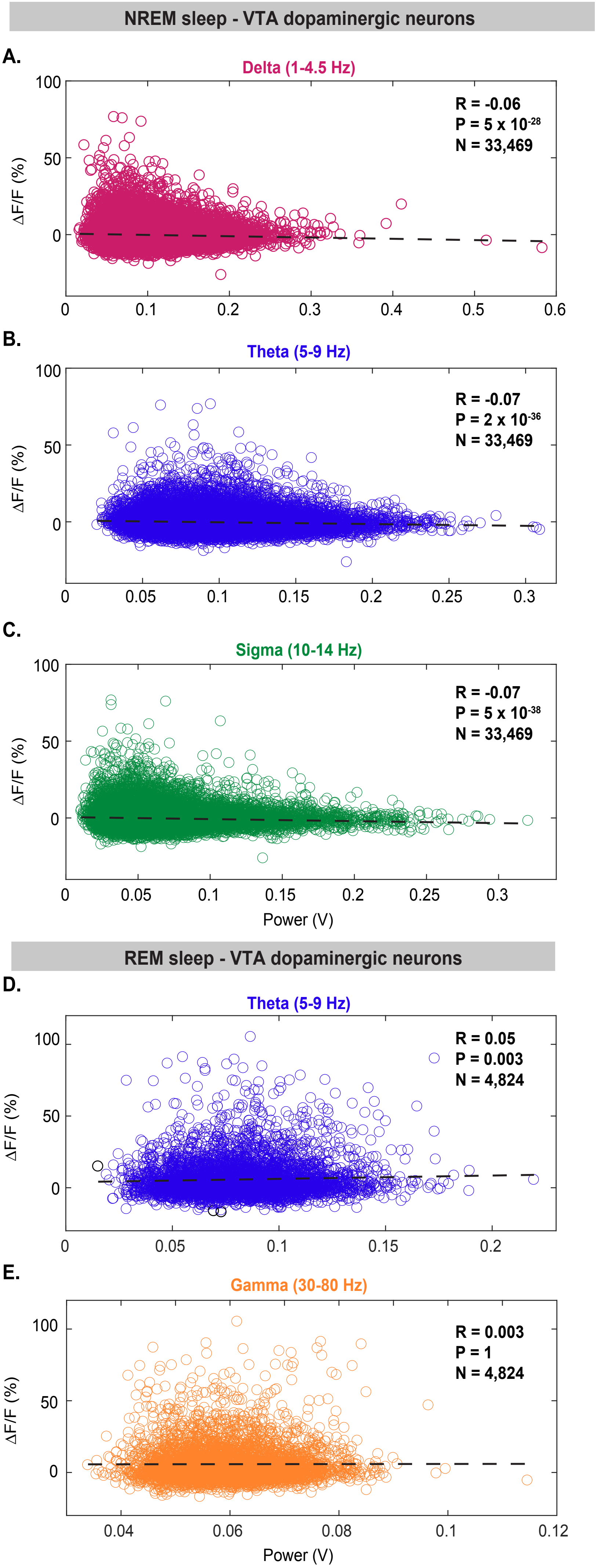
